# Physiological and injury-induced microglial dynamics across the lifespan

**DOI:** 10.1101/2024.10.02.615212

**Authors:** Taryn Tieu, Anne-Jolene N. Cruz, Jonathan R. Weinstein, Andy Y. Shih, Vanessa Coelho-Santos

## Abstract

Microglia are brain’s resident immune cells known for their dynamic responses to tissue and vascular injury. Little is known about how microglial activity differs across the life-stages of early development, adulthood, and aging. Using two-photon live imaging, we show that microglia in the adult cerebral cortex exhibit highly ramified processes and relatively immobile somata under basal conditions. Their responses to injury occur over minutes and are highly coordinated neighboring microglia. In neonates, microglia are denser and more mobile but less morphologically complex. Their responses to focal laser-induced injuries of capillaries or parenchymal tissue are uncoordinated, delayed and persistent over days. In the aged brain, microglia somata remain immobile under basal conditions but their processes become less ramified. Their responses to focal injuries remain coordinated but are slower and less sensitive. These studies confirm that microglia undergo significant changes in their morphology, distribution, dynamics and response to injury across the lifespan.

**HIGHLIGHTS:**

- Microglia undergo significant morphological and dynamic changes between life-stages.
- Neonatal microglia are highly dynamic, less morphologically complex and mount delayed responses to focal injury compared to adult microglia.
- Aged microglia are slightly less ramified and their responses to focal injury are slow and less sensitive than adult microglial.
- Maturation of microglial morphology in the developing cortex is disrupted by focal laser injury.

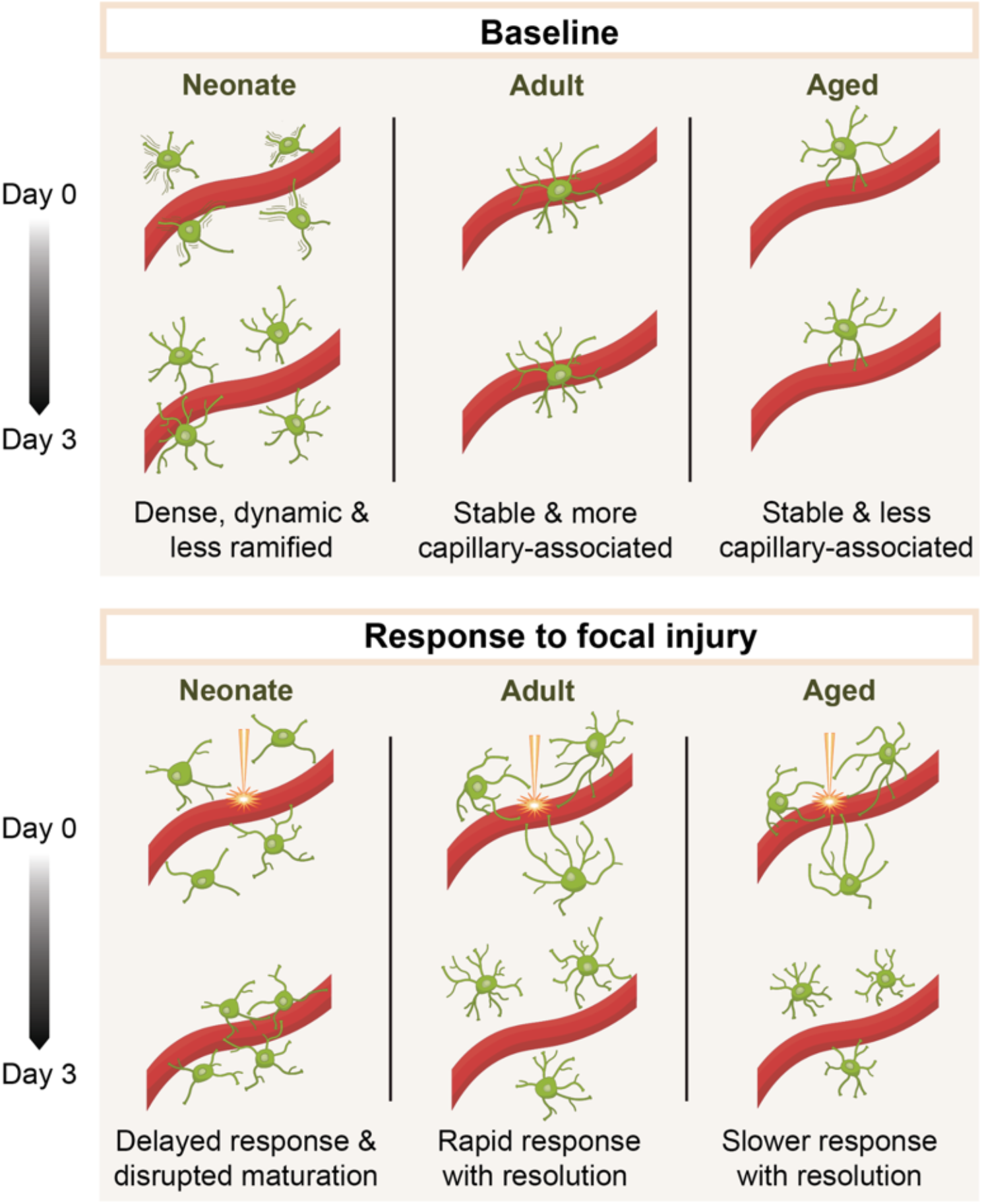

## INTRODUCTION

Microglia are resident immune cells of the brain, uniquely capable of modifying their morphology on a rapid time-scale of minutes while serving diverse roles in neurodevelopment, immune surveillance, and response to tissue injury^1^. Originating from the monocyte/macrophage lineage, microglial progenitors arise from meningeal macrophages and mesenchymal tissue to colonize the central nervous system during embryonic and early postnatal periods, prior to closure of the blood brain barrier (BBB)^2^. During development, microglia help to shape neural circuitry by pruning excess neuronal synapses and facilitating cell differentiation and maturation^3,4^. In adulthood, microglia continue to shape neural connectivity by modifying synapses with changes in sensory experience^5^.

In the first postnatal week of murine brain development, microglia exhibit an amoeboid shape and limited arborization of their processes. They bear morphological similarity to macrophages, and exhibit phagocytic roles for removal of cellular debris during shaping of the brain architecture^6^. Further, observations of physical associations between microglia and the vasculature in the developing brain suggest that they may support the development of microvascular structure and function, and/or use blood vessels to migrate into the brain^7,8^. Over the following weeks, microglia then undergo a morphologic shift from an ameboid shape to mature resting microglia, characterized by small somata and thin and highly branched processes extending radially from the soma^9^.

In the adult brain, microglia aid in maintenance of cerebral homeostasis and regulation of the neural environment^10–12^. As part of the brain’s innate immune system, microglia recognize injuries or pathogens and quickly initiate an inflammatory response^13,14^. They rapidly extend their processes and migrate towards the site of injury to act as a barrier between the injured and healthy tissue^15,16^. Microglia also re-acquire a developmental-like amoeboid morphology to change from resting to activated states in response to injury^17^. A well-coordinated immune response by microglia can have beneficial effects in the wound healing process, by contributing to clearance of cellular debris, neuroprotection, and limited extravasation of blood molecules from damaged vessels^18^. However, activation of microglia can be both beneficial or harmful depending upon the type of stress and damage signals, duration of injury, the tissue microenvironment, and the age of the organism^19^.

With brain aging, microglia become less ramified^20,21^. They increase the expression of genes involved in neuroprotection with brain aging, suggesting a compensatory response to concomitant age-related degenerative processes^22,23^. However, microglia may also become senescent or dystrophic, adopting dampened neuroprotective function with elevated production and release of pro-inflammatory cytokines and decreased production of anti-inflammatory cytokines, that can contribute to a chronic inflammatory state, neuronal dysfunction and cell death ^21,24,25 26,27^.

While many studies have used in vivo imaging to characterize microglial dynamics in the adult mouse brain, there remain major gaps in how microglia behave across life stages *in vivo*. To address this issue, we have used *in vivo* two-photon imaging through thinned–skull cranial windows to study microglial dynamics in the mouse cerebral cortex during development, adulthood, and in aging. We study their morphologies and dynamics under basal, resting conditions and in response to focal laser-induced injuries.

## RESULTS

### Distribution and morphological characterization of microglia across age

To visualize microglial morphology *in vivo*, we created thinned skull cranial windows over the somatosensory cortex in CX3CR1-GFP^+/-^ mice^15,16^ at three age periods: Neonatal (P9-12), adult (P90-155), and aged (P650-713). Thinned skull windows were used to minimize neuroinflammatory responses induced by cranial window surgery^28,29^. The cortex was imaged using two-photon microscopy to capture signals from GFP+ microglia (green) and the vasculature (red), which was labeled by intravenous injection of a high molecular weight fluorescent dextran dye (**Fig. 1A**). Image stacks were collected typically 90 µm from the pial surface up to 120 µm in depth.

**Figure 1.**
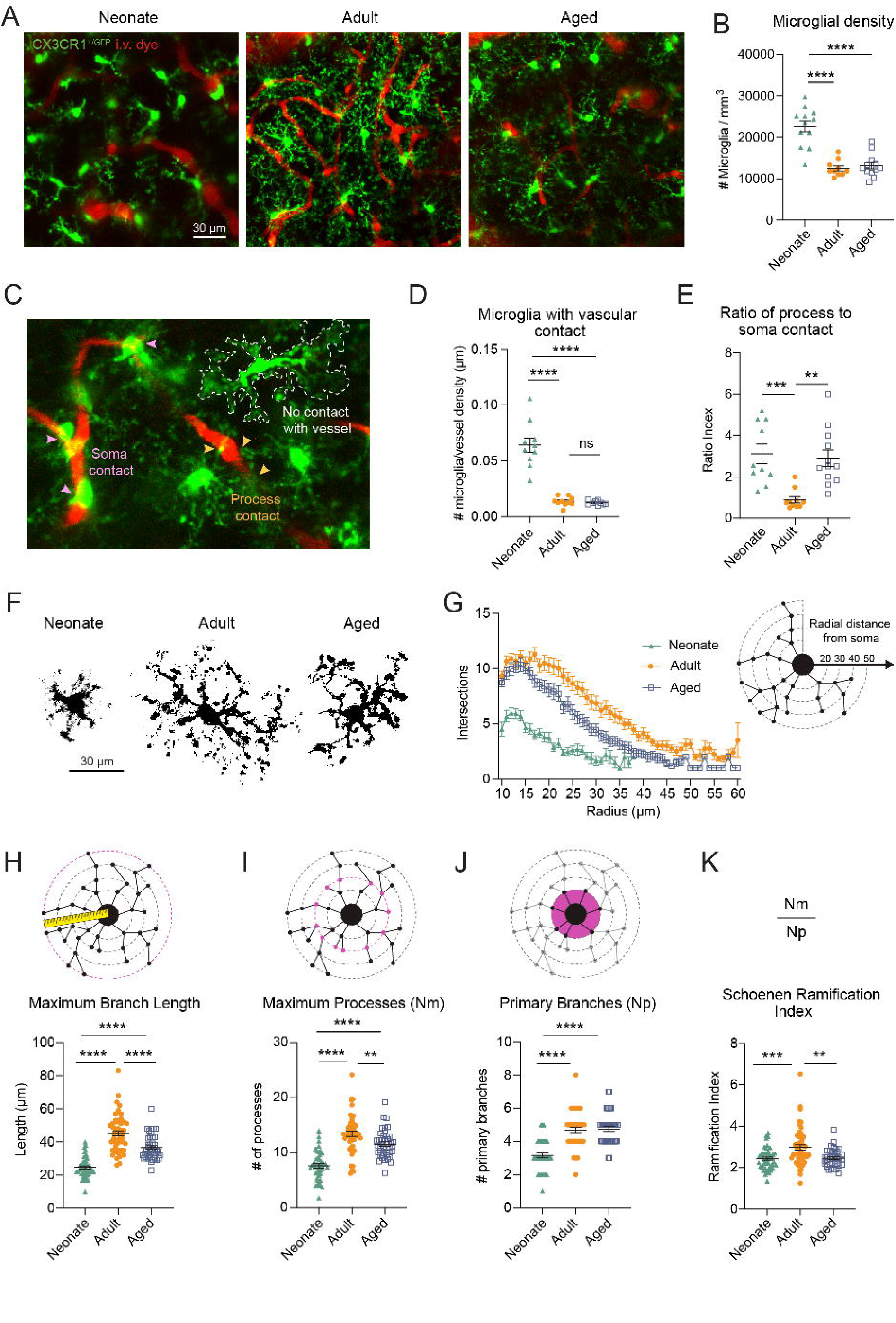
Physiological characteristics of microglia distribution and branch complexity. **(A)** Microglia distribution in neonates (postnatal 9), adults (3-5 months), and aged mice (21-23 months). **(B)** Quantification of microglia density. (n=12 fields from 8 neonatal mice, n=10 fields from 6 adult mice, n=12 fields from 5 aged mice) ****p<0.0001 using One-Way ANOVA with Tukey’s multiple comparisons test. **(C)** Examples of microglia soma, process and no contact with vessel. **(D)** Percentage of microglia contacting the vasculature via their processes or soma. (n= 11 fields from 8 neonatal mice, n=10 fields from 6 adult mice, n=12 fields from 5 aged mice, ****p<0.0001 using One-Way ANOVA with Tukey’s multiple comparisons test). **(E)** Ratio index of process-to-soma contact per field of view (n= 11 fields from 8 neonatal mice, n=10 fields from 6 adult mice, n=12 fields from 5 aged mice, ***p=0.0009 (neonatal vs adult) and **p=0.0016 (neonatal vs aged) using One-Way ANOVA with Tukey’s multiple comparisons test). **(F)** High magnification of a single microglia to demonstrate differences in morphology between neonate, adults, and aged. Scale bar 30µm **(G)** Result of Sholl analysis method used to map out processes and determine ramification index. Data are presented as mean ± SEM, (n = 45 cells from 4 neonatal mice, n=51 cells from 6 adult mice, n= 40 cells from 5 aged mice) One-Way ANOVA with Tukey’s multiple comparisons test. **(H-K)** Comparison of Sholl analysis data in postnatal, adult, and aged mice **(H)** Length from soma to the longest process point. **(I)** Highest number of process intersections that are occurring at one radius point. **(J)** Number of initial processes stemming from the soma. **(K**) Ratio of the process maximum to the primary branches, i.e., Schoenen Ramification index.

An initial survey of microglial glial density in image stacks revealed that microglia in the developing cortex were ∼2-fold higher than that seen in the adult and aged brain (**Fig. 1B** and **Supp. Fig. 1A**). Further, microglial density did not differ between adult and aged stages, suggesting establishment of a steady state distribution^30^. Recent work has revealed that microglial somata can make direct contacts with the endothelium^31^, dependent on their proximity to the vessel wall, and these interactions may contribute to blood flow regulation^32–34^.To explore this spatial heterogeneity, we evaluated vessel contacts (**Fig. 1C**) by distinguishing between juxtavascular microglia (soma or processes in contact)^35^ and microglia with no obvious contact. These analyses did not distinguish vascular zone-specific contacts, such as capillary-associated microglia (CAMs)^32^, as the capillary zone is still being established in the neonatal stage making comparisons across age more difficult^36^ (**Supp. Fig. 2**). Interestingly, neonates exhibited more microglia in contact with vessels (**Fig. 1D**), despite having less overall vascular density (**Supp. Fig. 2**). To more closely examine soma contacts that may infer direct microglial interaction with the endothelium, we also calculated the ratio of microglial process versussoma contact (**Fig. 1E**). This analysis revealed a significantly higher chance of soma-vessel contact in adults.

**Figure 2.**
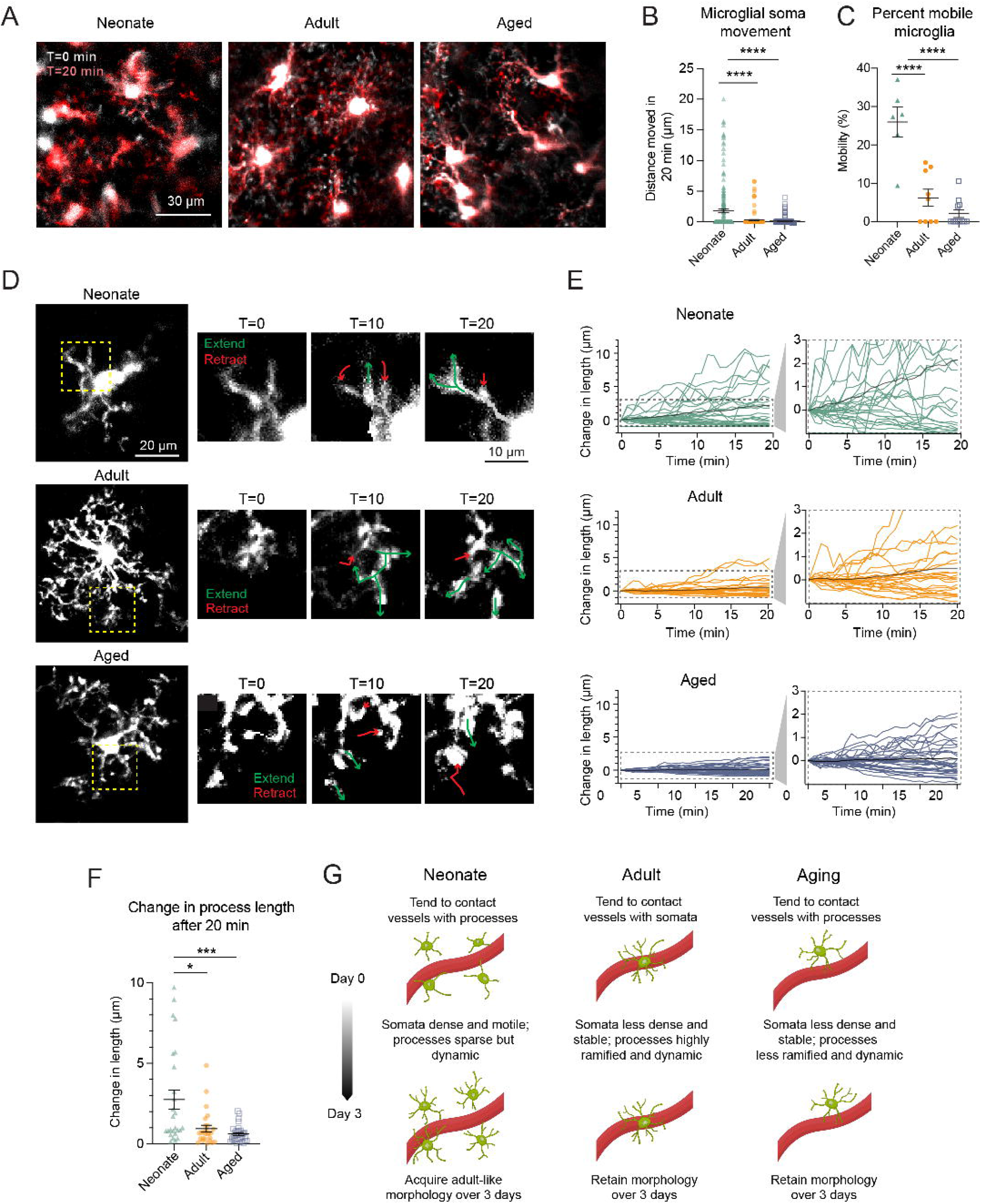
Microglia soma and process dynamics through the life stages. **(A)** Representative images showing microglia movement over 20 minutes in all ages. Light gray microglia represent the initial time of imaging, and it is overlaid with red colored microglia that depicts 20 minutes later. **(B)** Distance traveled by microglia somata. (n=195 cells from 4 P9 mice, n=144 cells from 4 adult mice, n=204 cells from 3 aged mice) ****p<0.0001 using One-Way ANOVA with Kruskal-Wallis multiple comparisons test **(C)** Percentage of mobile microglia in all age groups (n=6 cells from 4 neonate, n=9 cells from 5 adult mice, n=12 cells from 5 aged mice) ****p<0.0001 using One-Way ANOVA with Tukey’s multiple comparisons test **(D)** Representative images of microglial process extension and retraction at 0 min, 10 min, and 20 min in all ages. **(E) L**ength changes of individual microglial process movements in neonate, adults, and aged animals (n=650 processes from 3 neonatal mice, n=650 processes from 3 adult mice, n=846 processes from 3 aged mice). **(F)** Comparison of the change in process length at 20 min in all ages. *p<0.0165, ***p<0.0004 using One-Way ANOVA with Kruskal-Wallis post hoc test. **(G)** Graphic summary of microglia association with vessels and mobility results.

We next assessed microglial morphology (**Supp. Fig. 1B**) across ages using Sholl analyses of maximally projected 2D images (**Fig. 1F-K**). Across all ages, the density of microglial processes was highest close to the soma and then steadily decreased with greater distance from the soma (**Fig. 1G**). The processes of adult microglia extended the furthest from the soma, and exhibited the greatest density of branches. Unlike adult and aged microglia, neonatal microglia extended about half the number of processes, and the processes reached approximately half the radius of adult and aged microglia. Aged microglia were morphologically more similar to adult microglia, but exhibited slightly lower process density further from the soma (**Fig. 1H**).

Sholl analysis also reports the highest number of processes at an intersection (maximum processes) and the number of processes stemming directly from the microglial soma (primary branches). There was no significant difference between the number of primary branches in adult and aged microglia, both averaging around 5 initial processes. However as they ramified, aged microglia exhibited fewer processes than adults, as shown by maximum process values (**Fig.1 I-J**). In neonates, the primary branch number is lower at ∼3 initial branches and we also detected a notable decrease in the maximum process compared to the adult and aged groups (**Fig.1 I-J**). The Schoenen ramification index is a ratio of the maximum processes and primary branches, which provides a metric of process complexity. The ramification index was higher in adults compared to both neonates and aged mice, and there was no difference when comparing neonate and aged microglia (**Fig. 1K**). Thus, there are notable shifts in microglial morphology and the complexity of their processes between life stages.

### Microglial surveillance and dynamics across age

Live imaging enabled tracking of microglial dynamics (**Fig. 2A**). Over periods of 20 minutes, we found that microglia in neonates traveled a greater distance through the parenchyma compared to those in adult and aged groups (**Fig. 2B,C**). Adult and aged microglia exhibited small shifts in soma position but were overall quite stable in position over the 20 min time-frame. Microglia survey their microenvironments with dynamic processes, and this feature was also analyzed (**Fig. 2D**). Microglia in neonates exhibited larger and more rapid shifts in process length, which could be either extensions or retractions, in comparison to adult microglia (**Fig. 2E**). Changes in process length of neonatal microglia was significantly greater than adults and aged microglia (**Fig. 2F**). A trend toward a small reduction in process dynamics was observed in aged microglia compared to adult microglia (**Fig. 2F**). Note that our analysis of process length change examined growing processes that exhibited more extension than retraction of protrusions and was biased toward process extension. (**Fig. 2F**). In brief, the most dynamic microglia are observed in the neonatal brain both with respect to soma migration and processes surveillance (**Fig. 2G**).

### Microglia response after a focal laser injury: parenchyma *vs* vascular

Laser-induced focal injuries are commonly used to evaluate microglia response to injury during imaging^16,37^. We used this method to compare microglial responses to parenchyma and microvascular injuries across ages. Laser injuries to the brain parenchyma cause release of ATP, which induces the chemotactic response of microglial processes^16^, while microvascular injury involves further leakage of blood products known to induce clustering of microglia, such as fibrinogen^38^. We first examine responses to parenchymal injury (**Fig. 3A-C**), by measuring change in GFP intensity within a region-of-interest (ROI) demarcating the injured site where processes of neighboring cells converge (**Fig. 3A, B**). This reveals that adult microglia mounted a rapid and coordinated response, while aged microglia significantly reduced in their response, consistent with prior imaging studies^39^ (**Fig. 3C**). Further, we observed that neonatal microglia, similar to the aged brain, lacked a response to parenchyma injury (**Fig. 3C**). In both neonates and aged mice, microglial processes largely fail to reach the parenchymal injury region within 16 min, in contrast to that seen in adults (**Supp. Fig. 4**). All mice were re-examined 3 days after injury to understand sub-acute microglial responses. In adults, GFP intensity in the ROI returned to baseline within 3 days, while aged mice showed a persistent microglial aggregation, and neonates exhibited an even further increase of microglial response (**Fig. 3C**).

**Figure 3.**
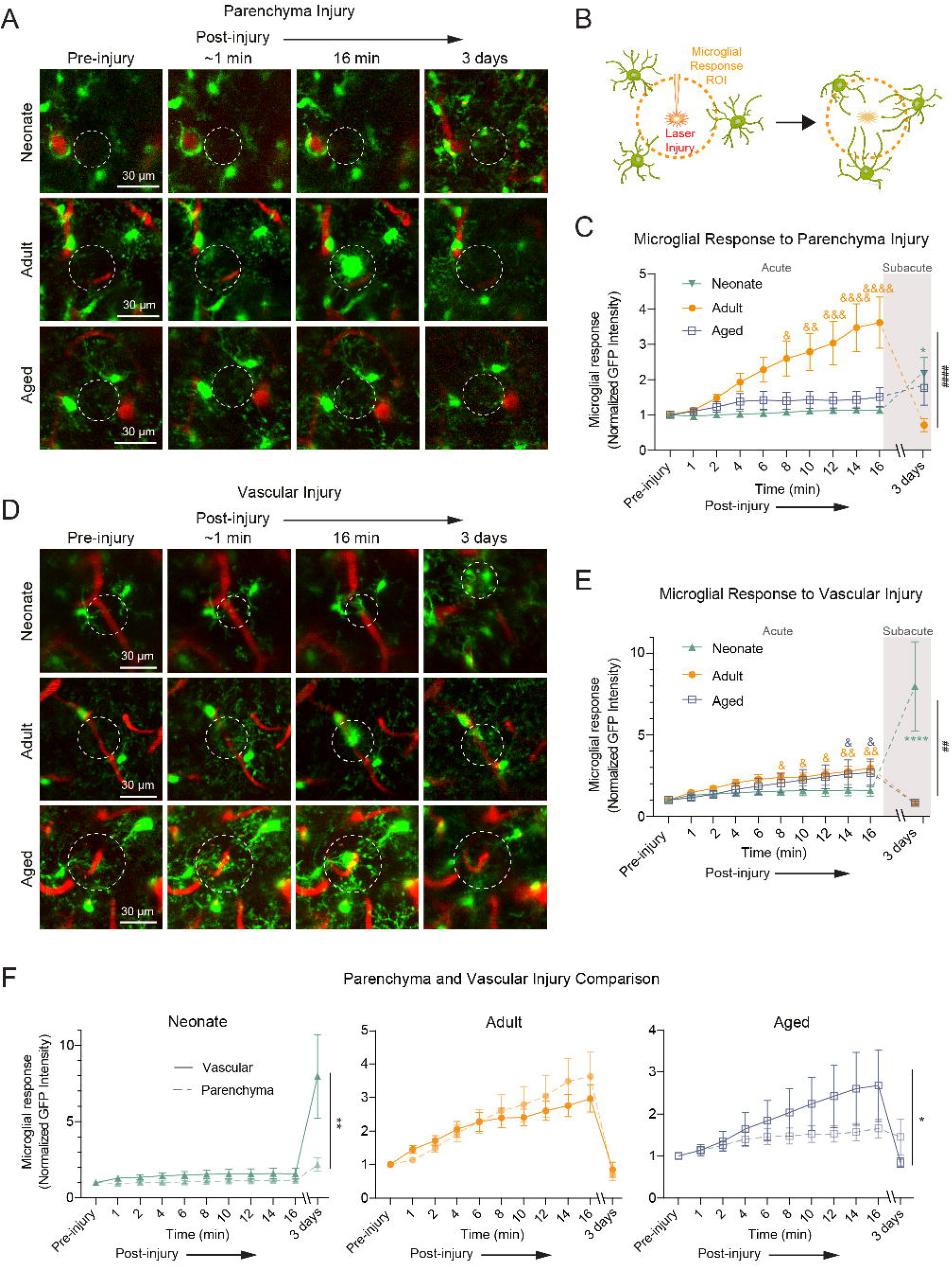
Comparing microglia reactivity in parenchyma and vascular injuries. **(A)** Representative images of microglia reaction to parenchyma injury. The mice were reimaged 3 days later. **(B)** Schematic of laser injury experiment. Region of interest is a demarcated area around the injury site where the microglia response is measured. **(C)** Normalized fluorescence intensity of parenchyma injury sites collected to measure microglial response. (n=14 areas from 6 neonatal mice, n=11 areas from 8 adult mice, n=8 areas from 4 aged mice). Acute response (0 to 16 min): &p=0.0228; &&p=0.0057; &&&p=0.0007 and &&&&p<0.0001 using Two-way ANOVA with Sidak’s post hoc test, only adults respond to the injury. Subacute response (pre-injury vs 3 days): *p=0.0134 using Two-way ANOVA with Dunnett’s post hoc test, only neonatal was impacted. Overall response to parenchymal injury: ####p<0.0001 neonate vs. adults and aging vs. adults using Two-way ANOVA with Dunnett’s post hoc test. **(D)** Representative images of microglia reaction to vascular injury. The mice were reimaged 3 days later. **(E)** Normalized fluorescence intensity of vascular injury sites collected to measure microglial response. Data are presented as mean ± SEM, (n=14 areas from 6 neonatal mice, n=11 areas from 8 adult mice, n=8 areas from 4 aged mice). Acute response (1 to 16 min): adults pre-injury vs acute 8 min &p=0.0307, 10 min &p=0.0281, 12 min &&p=0.0071, 14 min &&p=0.0021, 16 min &&p=0.0004; aging pre-injury vs acute 14 min &p=0.0365, 16 min &p=0.0238 and Two-Way ANOVA with Sidak’s post hoc test. Subacute response (pre-injury vs 3 days): ****p<0.0001 using Two-way ANOVA and Dunnett’s post hoc test, only neonatal was impacted. Overall response to parenchymal injury: ##p=0.0044 neonate vs. adults using Two-way ANOVA with Dunnett’s post hoc test. **(F)** Comparison of parenchyma and vascular injury response in each neonate, adult, and aged groups. (n=14 areas from 6 neonatal mice, n=11 areas from 8 adult mice, n=8 areas from 4 aged mice) *p=0.0151 and **p=0.0010 parenchyma vs vascular injury using Two-way ANOVA with Sidak’s post hoc test. Scale bar 30µm

**Figure 4.**
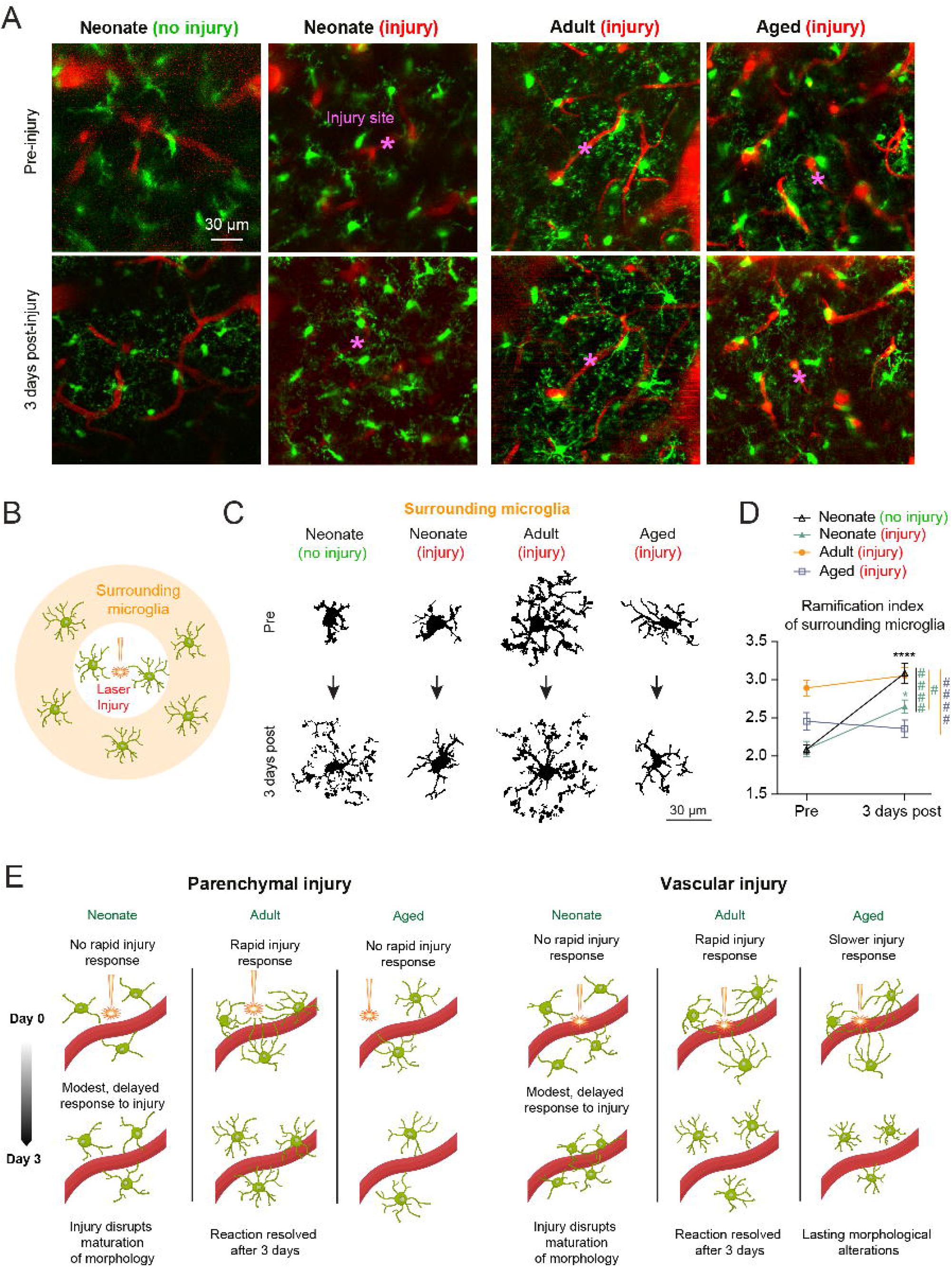
Assessing peripheral microglia response to injury. **(A)** Full-field of view representation at 0 minutes and 3 days of neonates with no injury and neonates, adult and aged mice with vascular injuries. Asterisks mark where the injury was made. Scale bar 30µm. **(B)** Schematic of surrounding microglia, defined as microglia in the imaging field outside of the Injury ROI. **(C)** Comparison of individual surrounding microglia examples in each experimental group at 0 minutes and 3 days after vascular injury. **(D)** Ramification index of surrounding microglia in vascular injury experiments (n=14 areas from 6 neonatal mice, n=11 areas from 8 adult mice, n=8 areas from 4 aged mice). Overall response to vascular injury: ****p<0.0001 neonate non-injury and *p=0.0309 neonate non-injury using Two-way ANOVA with Sidak’s post hoc test. Subacute response (pre-injury vs 3 days): ####p<0.0001 neonatal injury vs. non-injury and adults vs. aged; #p=0.0208 neonatal injury vs. adults using Two-way ANOVA with Tukey’s post hoc test, only neonatal was impacted. **(E)** Graphic summary of parenchyma and vascular injury response.

Microglial responses to vascular injury were also assessed in all ages (**Fig.3D**). While vascular injury likely led to some extravasation, the use of high molecular weight dextran (2MDa) precluded visualization of mild blood leakage. The stalling of blood flow (loss of capillary flow) at the injury site was used as an index of sufficient injury. Interestingly, blood flow stalling was resolved within 16 min in most injured vessels of adult and aged mice, while many vessels in neonates remained stalled over the acute observation period. However, blood flow was eventually regained at the 3 days post-injury time-point (**Supp. Fig. 3**). Interestingly, while adult and aged microglia showed a coordinated acute response to laser induced injury, neonatal microglia largely lacked this response. When mice were re-examined 3 days later, the adult and aged microglial response returned to baseline level of GFP fluorescence, while neonates significantly increased, again suggesting that there is a delayed response to vascular injury during development (**Fig. 3E, Supp. Fig.5**).

We further compared both injury types within each age group and found that the adult microglia behaved similarly between these two injury types, while aged microglia showed more modest responses to parenchyma injuries compared to vascular injuries. In neonates, microglia exhibited similar minimal responses to both injury types but pronounced responses at 3 days with greater reaction to vascular injury (**Fig. 3F**).

### Focal injury inhibits morphological development of microglia in neonates

Brain injury can have an enduring effect on microglial activation state in tissues well beyond the injury site^40^. We therefore examined how microglia surrounding the focal injury to a microvessel behaved over several days post-injury (**Fig.4 A-B**). These microglia may be several hundreds of micrometers to a millimeter away from the injury, but within the same thinned-skull window. We compared morphology pre-injury and 3 days post-injury for all age groups (**Fig.4 C**). Further, an additional control group with no laser injury was also examined to understand normal developmental shifts in microglial morphology. This revealed that from P9 to P12 of normal development, microglia transitioned from an ameboid morphology into a highly ramified morphology moving toward the complexity of those seen in adult cortex. However, microglia surrounding the injury site were significantly less ramified by P12 and retained an ameboid morphology (**Fig. 4D**). We also performed this analysis for the parenchyma injury group, and found a similar block in the shift from ameboid to ramified morphology 3 days post-injury, although it was less pronounced than with vascular injury (**Supp. Fig. 6**). In the adult and aged brain, surrounding microglia exhibited little morphological change 3 days post-injury.

Thus, both tissue and vascular injury may delay the maturation of surrounding microglia in the developing brain.

## DISCUSSION

In this study, we addressed a key gap in knowledge on how microglial behaviors change across different life stages. We characterized microglial morphology and dynamics in both healthy conditions and following focal, laser-induced injury in neonatal, adult and aged mice (**Fig. 4E**). These experiments offered a unique look into the neonatal brain and revealed marked differences between developing and adult microglia. They also provided insight on how microglia, known to be long-lived, differ in their behaviors with brain aging. Across the lifespan, we identified distinct physiological characteristics and responses to two forms of tissue injury.

Neonatal microglia exhibited the greatest dynamics, both in terms of soma and process mobility. These enhanced dynamics may help compensate for reduced structural complexity, and enable broader parenchymal coverage for effective tissue surveillance. Increased dynamics may also be associated with microglial migration. Prior studies have demonstrated that juxtavascular microglia are more prevalent during early postnatal stages (P1-5), as they migrate along blood vessels to colonize the brain^35^. However, by P7-14 and continuing into P21, their prevalence drops to less than 20% of the total microglia, as they move to proliferate in the tissue parenchyma. Consistent with these findings, our results also showed greater juxtavascular microglia in neonates and a decrease in adulthood. The relation between an “ameboid” morphology of neonatal microglia and its role in brain physiology remains limited as emerging evidence suggests that microglial morphology is complex and cannot be simply defined as “activated”^41,42^. However, the amoeboid shape of neonatal microglia may also indicate a phagocytic phenotype^43^. Through phagocytosis, these cells play a vital developmental role in refinement of neuronal circuits by maintaining the neuronal population, eliminating dead cells, and pruning synapses^44^.

Microglia in aged mice exhibited low process complexity similar to the neonate, but displayed a different pattern of behavior in vivo. Their somata remained stationary and their process dynamics were slower, leading to diminished surveillance capabilities^45^. Aged microglia are associated with amplified pro-inflammatory responses, resulting in a chronic inflammatory state often referred to as “primed” microglia^46^. This persistent inflammation may contribute to their low ramification index, even in the absence of overt pathology^47^. However, some evidence suggests that these cytoplasmic changes might not be solely indicative of inflammation but may also reflect characteristics of senescence and dystrophy^48^.

A molecular basis for why neonatal and aging microglia have a lower ramification index than adults and react preferentially to vascular injuries remains unclear. However, some microglial receptors, such as transmembrane protein 119 (TMEM119) and purinergic receptor P2Y, G-protein coupled 12 (P2ry12) that are critical for the injury response, undergo changes in expression with age. In the neonatal brain, these receptors increase in expression but might not be fully operational due to ongoing maturation^49^. In aging microglia, their downregulation or dysfunction seems to occur^50^.

In neonates, there is a delayed reaction to focal injury, and persistent reduction in branching complexity during a timeframe when microglia transition from ameboid to highly-branched surveillant cells of the adult brain^51^. These slower and persistent microglial responses stand in stark contrast to rapidly resolved responses in adult and aged mice, suggesting a lasting influence on the developmental process. In prior studies, Raghupathi et al. examined microglial responses to single and repetitive injuries in neonatal rats, particularly in the context of abusive head trauma. They observed increased microglial activity from 1 to 40 days post-injury, with more pronounced effects in repetitively injured brains^52^. This was associated with cognitive deficits, traumatic axonal injury, neurodegeneration, and sustained spatial learning deficits within two weeks following multiple impacts. Disruptions in neonatal microglia caused by systemic or central inflammation, traumatic brain injuries, or abusive head trauma can lead to significant neurological consequences^53^. Neuroinflammation has been linked to an increase in neurodevelopmental disorders later in life^54,55^. For example, autism spectrum disorder (ASD) has been associated with ongoing elevated immune responses and inflammatory phenotypes, including increased expression of MHC class II and inflammatory factors^56^, alongside morphological changes in microglia such as soma enlargement and process retraction^57^. Similar inflammatory responses and microglial activation surrounding vasculature were observed by us in attention deficit hyperactivity disorder (ADHD)^58^, with findings indicating that activated microglia contribute to these conditions by damaging the endothelial barrier. More recently, it was shown that microglial activation in early postnatal development prevents the normal pruning of cerebellar neurons, resulting in a permanent increase in cerebellar granule neurons and ADHD-like behaviors^59^. These findings emphasize the importance of maintaining proper microglial function during development to prevent long-term neurodevelopmental disorders^60^.

Another intriguing result was that aging microglia preferentially reacted to vascular damage. Compared to the adult, the response to vascular injury was preserved, while response to parenchymal injury was diminished with age. It is possible that the vascular injury produced in aged mice was more severe due to aged-related degradation of vascular structure^61^. Indeed, blood flow in some of the injured vessels did not fully recover, suggesting that the aged vasculature is more vulnerable. Additionally, aging microglia exhibit reduced soma contact with blood vessels. How this shift factors into age-related blood flow changes and regulation of capillary tone warrants further studies^32,33,62^.

## LIMITATIONS OF THE STUDY

The short study period may limit the generalizability of findings over longer developmental phases, as the study design was unable to assess long-term effects of injury and microglial changes. Further, the studies were conducted on mice missing one allele of CX3CR1, and prior studies have shown that this can dampen the inflammatory response^63^, though not as severely as loss of both alleles^64^. Further, the developmental effects of CX3CR1 haploinsufficiency are also not well understood. Additionally, the lack of sex-based analysis leaves potential differences unexplored. Despite these limitations, our research lays the groundwork for future studies to further investigate these areas and enhance our understanding of microglial dynamics across life stages.

## Abbreviations

BBB: Blood-Brain Barrier
CAMs: Capillary-Associated Microglia
i.p.: intraperitoneal
i.v.: intravenous
GFP: Green fluorescent protein
ROI: Region Of Interest

## AUTHOR CONTRIBUTION

Experiments were designed by VCS and AYS. Experiments were conducted by TT, VCS and AYS. Data analysis was performed by TT, AJC and VCS. Statistics were performed by VCS. The manuscript was written by TT with contributions from JRW, AYS, VCS.

## ACKNOWLEDGMENTS

We thank Tiago Figueiredo for creating artwork used in the graphical abstract and the schematics on Figures 1 to 4 (www.behance.net/TiagoFigueiredoGD).

## Declaration of interests

The authors declare no competing interests.

## ONLINE METHODS

### Animals

The Institutional Animal Care and Use Committee at the Seattle Children’s Research Institute approved the procedures used in this study. The institutions have accreditation from AAALAC and all experiments were performed within its guidelines.

For this study we used CX3CR1-GFP/+ mice were obtained from Jackson laboratory (B6.129P2(Cg)-*Cx3cr1tm1Litt*/J; Jax ID: 005582) and only heterozygous mice were used for the study. We considered the day of birth as postnatal day zero (P0), and neonatal pups were between P9 and P12 when used. Adult mice were 3-5 months old, and aged mice were 21-23 months old.For all studies we used a roughly equal mixture of male and female offspring. The mice were housed on a 12-hour light (7am - 7pm) – dark cycle, with *ad libitum* access to chow and water. We did not employ randomized selection strategies for allocation of animals to different experimental groups. Experimental cohorts consisted of littermates as best matched as possible for age and sex. Due to limited availability and therefore sampling in postnatal and aged mice, we did not explicitly test for an effect of sex and therefore data was pooled across sex.

### Cranial window surgery

Anesthesia was induced with isoflurane (4% induction, 1-2.5% maintenance in medical-grade air. During surgery, mice are laid in a prone position on a feedback-regulated heat pad (FHC Inc.) to regulate body temperature at 37°C. To minimize disruption of the brain, thinned-skull cranial windows were generated in all animals as described previously^29,65^. The scalp was first cleaned with betadine and artificial cerebrospinal fluid (ACSF) before a midline incision was made starting between the eyes to immediately caudal of the ears. The windows were placed generally above the somatosensory cortex^66,67^ on the left hemisphere. Over the contralateral hemisphere, a head mount was affixed to the skull with cyanoacrylate glue (Loctite 401) and then Metabond dental cement was laid over the attachment as a reinforcement. This ensured steadiness when creating the thinned skull window and reduced movement artifacts during imaging. At this stage in neonatal surgery, buprenorphine (Patterson Veterinary) was provided at a dose of 0.05 mg/kg (postnatal) intraperitoneal injection (i.p.), for analgesia.

Cranial windows in adult and aged mice are performed while the skull was dry by carefully shaving away bone using a dental drill. In neonates, the cutting edge of a 31-gauge needle (BD Biosciences; catalog #305109) and a surgical scalpel blade (Fisher Scientific; #15; catalog #22-079-701) was used as their skulls were thinner and more delicate. Occasionally, the skull was moistened using ACSF to examine the window for translucency. Once the thinned skull was ready, the area was dried before a drop of cyanoacrylate glue (Loctite 401) was applied and a ∼3 mm diameter round coverslip for adults and aging or ∼1 mm precut square coverslip for neonates was laid over the thinned region. After the glue dried, dental cement was applied to the remaining uncovered areas around the window and a slight berm was created to hold water for the imaging lens. With surgeries on adult and aged mice, buprenorphine (Patterson Veterinary) was provided immediately post-surgery at a dose of 0.1 mg/kg, i.p., for analgesia. Imaging took place on the second day following surgery to allow for recovery.

### Two-photon microscopy

In vivo two-photon imaging system was performed with a Bruker Investigator (run by Prairie View 5.5 software) coupled to a Spectra-Physics Insight X3. Green and far red fluorescence emission was collected through 525/70 nm and 660/40 nm bandpass filters, respectively, and detected by GaAsP photomultiplier tubes (PMTs). High-resolution imaging was performed using a water immersion 20-X, 1.0 NA objective lens (Olympus XLUMPLFLN 20XW) using 975 nm excitation during the experiment. Lateral sampling resolution was 0.4μm per pixel and axial sampling was 1μm steps between frames. To label the vasculature, a custom conjugated Alexa Fluor 680-dextran^68^ (Alexa Fluor 680; Life Technologies; A20008 conjugated to 2 MDa Dextran; Fisher Scientific; NC1275021) was injected through the retro-orbital vein under deep isoflurane anesthesia. During imaging, isoflurane was maintained at 1.5% MAC in medical-grade air.

### Focal laser-induced injury

Before injury experiments began, mice were checked for sufficient water in the berm of the cranial window to ensure imaging quality throughout the entirety of irradiation and imaging process. A capillary-dense area at a depth of around 80-100*μ*m into the cortex was first located. In neonates there were non-flowing angiogenic sprouts, and we carefully selected connected, flowing vessels in order to mimic conditions across all ages. A single subset of pre-injury image stacks was then taken at 975nm with a depth of ∼20µm with 1µm steps. Microvascular injuries were targeted at the capillary wall and conducted at 800 nm excitation using a focused two-photon line-scan. A single vessel in the field of view was exposed to 60 seconds of focal irradiation at 100mW of power. Parenchymal injuries were targeted away from capillaries. Immediately after irradiation, the previous settings at 975nm are restored and the same 20µm thick images are taken over the course of 16 minutes to record post-injury microglial responses. Animals are re-imaged 3 days later to record longer-term post injury effects. Laser powers were determined at the output of the 20X objective using a laser power meter (ThorLabs; PM100D), with galvanometric mirrors engaged in a full field, 512 × 512 pixel scan covering 16µm ^2^. At 100 mW of power with a 60 second duration of scanning through a thinned skull, we found that only a minor injury was inflicted there was no overt rupture of targeted capillaries. Inducing injury at higher laser power induced larger reactions from the microglia which would obscure the ability to carefully observe microglial behavior.

### Microglia analysis

*Imaging and analysis of microglia soma movement:* ImageJ was used to measure microglial dynamics during basal physiological conditions. Using a 20 minute movie (5 frames per second 0.20s/frame using 3.6 μs/pixel dwell time), somata movement was assessed by calculating the difference in soma-vessel distance between an individual microglial soma to a nearby vessel from time 0 and time 20. Vessels remain stationary, thereby serving as a control against measurement errors due to any potential movement artifacts. We considered a cell mobile if it had traveled more than 2µm. Moving somata were counted and averaged to report the percentage of mobile microglia in each age group.

### Microglia process dynamics

To evaluate process dynamics, individual processes from multiple microglia were manually measured at every 2 minute time point and measured from their primary branching point on the soma to their most distant point. Processes were chosen based on best visibility. Any extended or retracted branching was also added or subtracted from the total length. **Fig 2 B to F** data was plotted as net change in process length with all individual values and also averages by age at each imaging time point.

### Sholl Analysis

A Fiji plug-in for ImageJ (http://fiji.sc/Sholl_Analysis) was used for branching patterns. Before starting the analyses, target cells were isolated and a default threshold adjustment was performed to produce a clearer image for the Sholl analysis to run. The threshold was set based on the best representation of each individual cell and its surrounding processes. The Z range in image stacks were adjusted to exclude any unwanted portions of neighboring cells or other background in the projected images used for analysis. Further, the drawing tool was used to carefully erase extra background noise. A straight line was manually drawn from the middle of the cell soma to the most distant process. The initial starting radius was based on the soma radius, which in our study was around 10µm. Neonatal microglia tended to have a higher radius due to a larger somata. Running the Sholl analysis created a series of concentric circles from the initial starting radius and was consecutively set to increase by 1µm until it reached the furthest process. The program then counted how many intersections were made with the processes of the cell at each circle. In order to analyze morphological characteristics, the raw data was processed to generate different sets of data: Maximum branch length (distance from the soma where the most branching intersections are occurring), maximum processes (the highest number of processes intersecting a radial region of interest), number of primary branches (how many initial branches extended from the soma), and a Schoenen ramification index (maximum processes / number of primary branches). We conducted Sholl analysis on data collected pre-injury, 16 minutes post-injury, and 3 days post-injury to assess microglial morphology in undisturbed, lesioned, and sham conditions. On the account of lack of in vivo research in microglial morphology in neonatal development, two-photon images were also taken of P12 neonates that had not been affected by injury experiments. Sholl analysis was also performed on these microglia and were plotted and compared with neonates with injuries at P9.

### Microglial density

Cell density per region was calculated as the number of microglia/volume in mm^2^, (196.61 × 196.61 × 33 µm). In addition, microglial cells were distinguished by interaction with the vasculature which was determined strictly by process or soma contact to blood vessels. Percentages of each type of interaction as well as no observable interactions were quantified from the total number of microglia per field. All density data are normalized to total vascular volume. Vascular length was analyzed using IMARIS version 10.1 through the 3D semi-automatic filament tracing method (auto-depth function).

### Microglia response

To measure microglial reaction to focal injury, fluorescence intensity over time was measured from ROIs using ImageJ/Fiji in pre-injury, post-injury (every 2 minutes), and 3 days post-injury images. The ROIs were determined by the space around the induced injury site, typically area around 500-1000*μ*m^2^ on average. ROIs size were occasionally adjusted to avoid high fluorescence signal contribution from microglia somata that could mask signal changes caused by process extension. Our data was then normalized to the first time-point within the movie, and then averaged by age groups. Additionally, randomized areas were chosen to measure fluorescence intensity of regions distant from the site of injury to assess peripheral microglial reactions.

### Exclusion criteria

Only microglia with all processes visible within the imaged area were included for either density, Sholl and dynamics analysis.

### Statistics

Statistical analyses were performed using Graphpad Prism 9 software. Statistical tests and details are provided in the Figure legends. Tests of normality were first performed to validate the use of parametric tests. When the assumptions of normality were not met, appropriate non-parametric tests were also employed.

## Notes

**Funding:** This work was supported by grants to A.Y.S. from the NIA and NINDS (NS106138, AG062738, AG081840, AG077731, NS097775), and to J.R.W from the NIA (NS124627). VCS was supported by the American Heart Association (20POST35160001) and by a Junior Leader Fellowship from the ‘La Caixa’ Foundation (LCF/BQ/PI22/11910036).

**Conflicts of interest:** The authors have no financial or non-financial conflicts of interest.

### Competing Interest Statement

The authors have declared no competing interest.

