## Supplementary Material for "Physiological and injury-induced microglial dynamics across the lifespan"

### Inventory of supporting information

**Supplementary Figure 1. Morphological characteristics of microglia across the lifespan.**

**Supplementary Figure 2. Vascular density in the somatosensory across the lifespan.**

**Supplementary Figure 3. Percentage of vessels regaining blood flow or stalled 16 minutes and 3 days post vascular injury.**

**Supplementary Figure 4. Proximity of microglia to site of parenchyma injury.**

**Supplementary Figure 5. Proximity of microglia to site of vascular injury.**

**Supplementary Figure 6. Assessing surround microglia response to injury.**

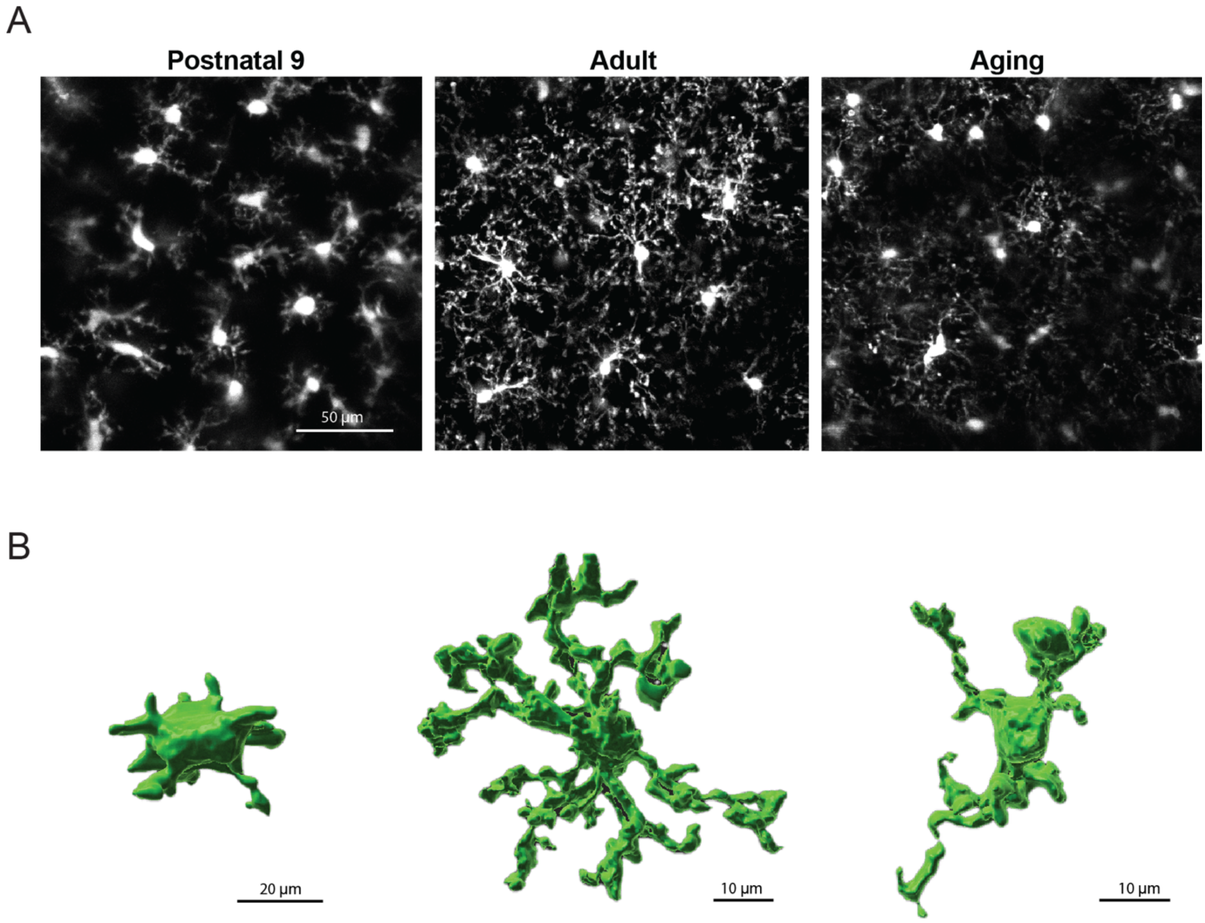


**Supplementary Figure 1. Morphological characteristics of microglia across the lifespan.** **(**A) Microglia distribution without blood vessels shown in neonates, adults, and aged mice. (B) 3D reconstruction of individual microglia representative of morphology in each age group.

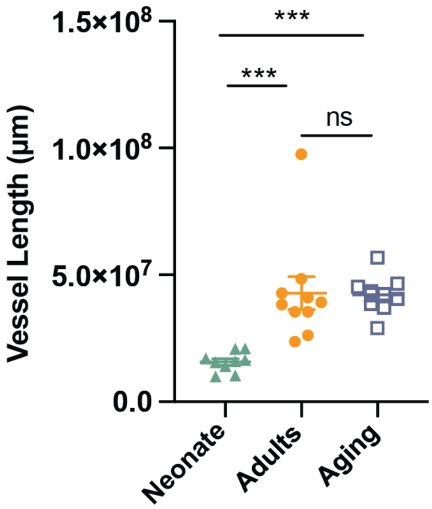


**Supplementary Figure 2. Vascular density in the somatosensory cortex across the lifespan**. From postnatal day 9 to adulthood, the vascular density doubles, indicating a significant increase in blood vessel formation during brain development. However, no significant differences in vascular density are observed in the cortical areas when comparing adulthood to aging, suggesting that the vascular structure remains relatively stable after reaching adulthood. Data are presented as mean ± SEM. n= 11 fields from 8 P9 mice, n=10 fields from 6 adult mice, n=12 fields from 5 aged mice, ***p=0.0004 (neonate vs adults) and ***p=0.0007 (neonate vs aging) using One-way ANOVA analysis with Tukey's multiple comparisons test.

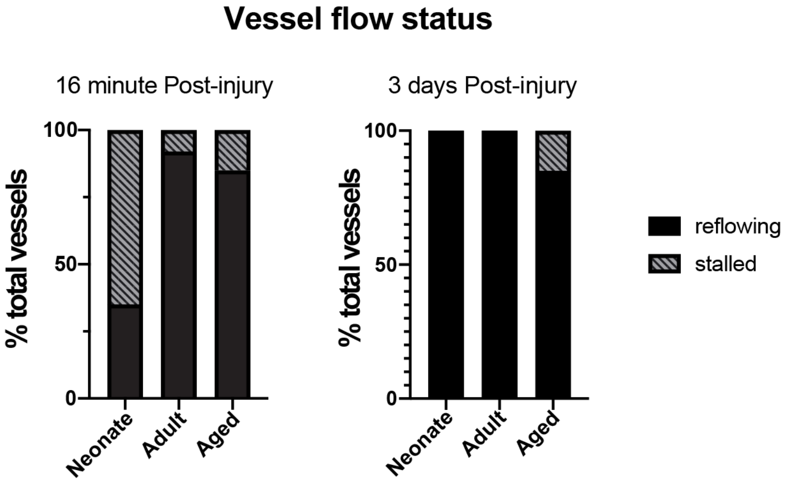


**Supplementary Figure 3. Percentage of vessels regaining blood flow or stalled 16 minutes and 3 days post vascular injury.**

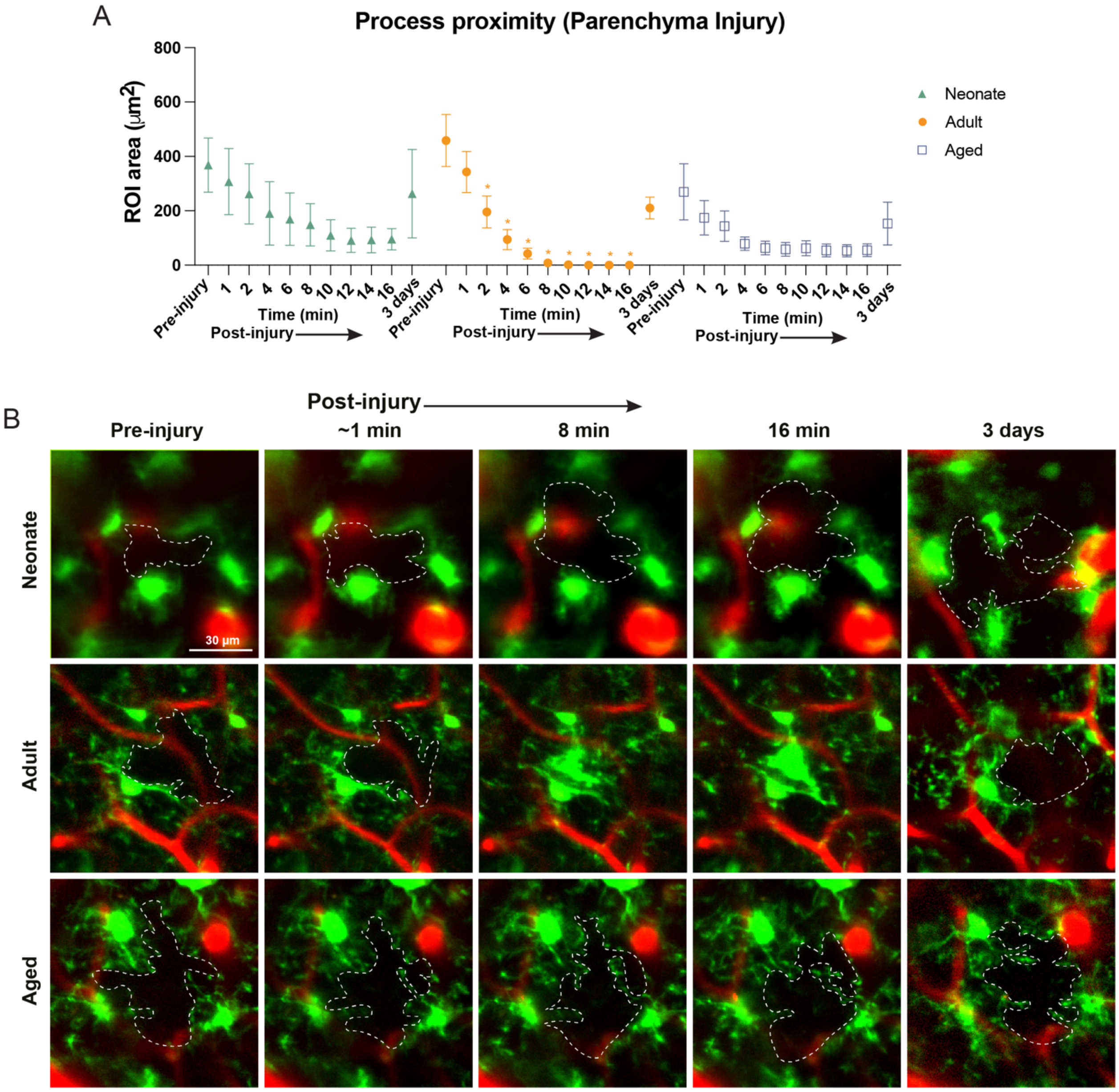


**Supplementary Figure 4. Proximity of microglia processes to site of parenchyma injury.** (A) Averages of the measured area between local microglia processes and parenchyma injury site. (B) Examples of areas around parenchyma injury site before injury and 1min, 8min, 16min, and 3 days after induced injury.

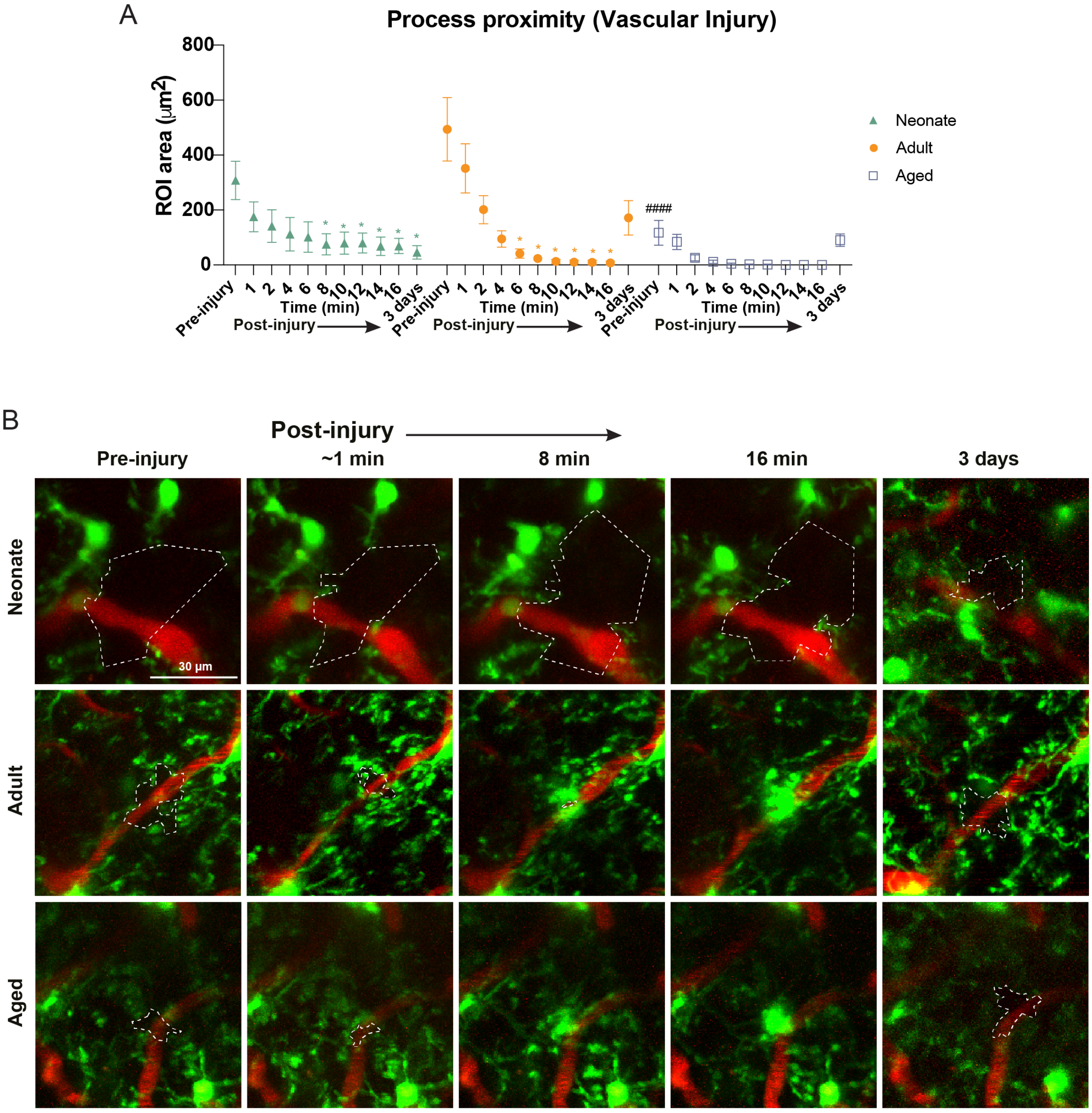


**Supplementary Figure 5. Proximity of microglia processes to site of vascular injury.** (A) Averages of the measured area between local microglia processes and vascular injury site. (B) Examples of areas around vascular injury site before injury and 1min, 8min, 16min, and 3 days after induced injury.

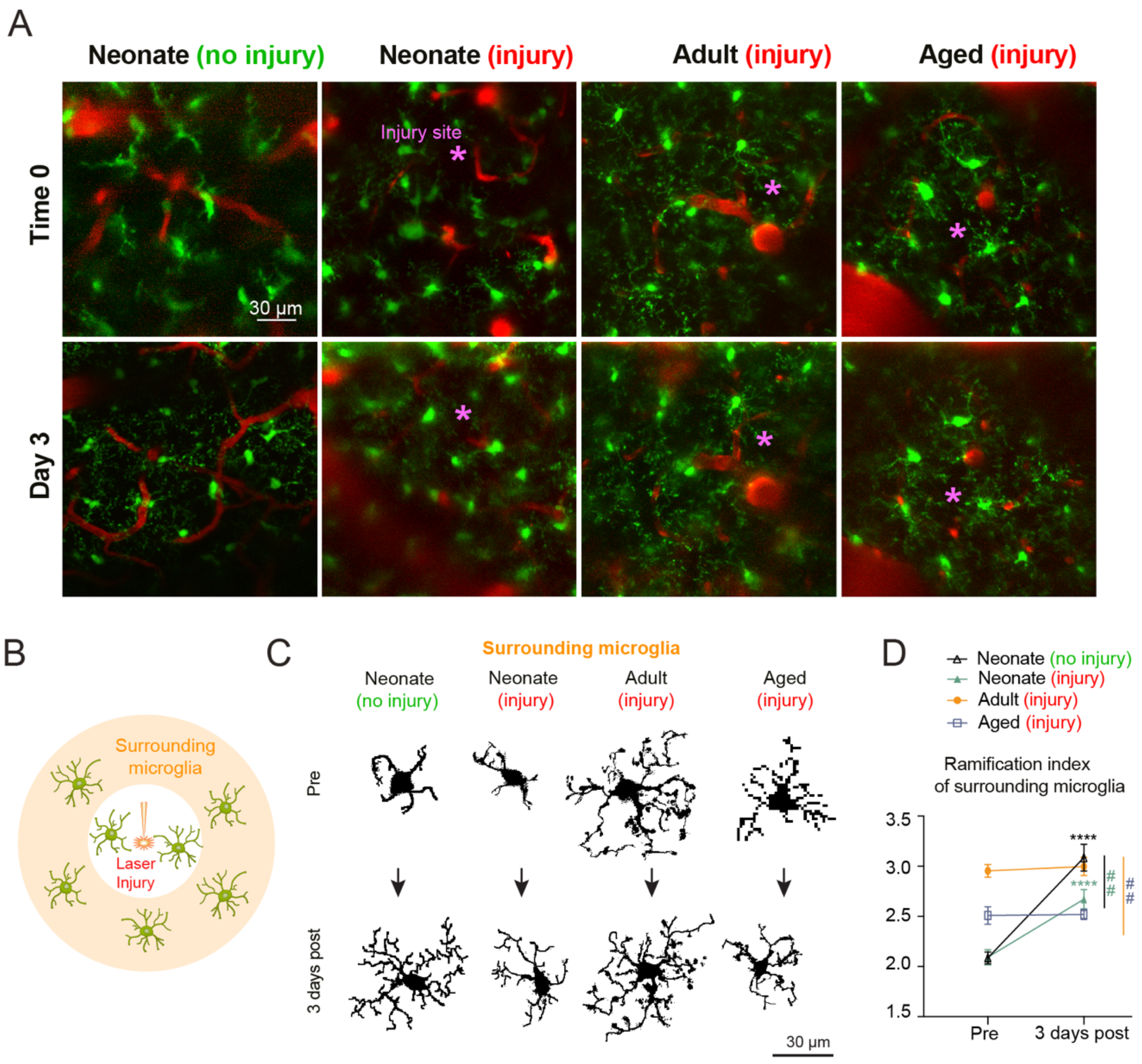


**Supplementary Figure 6. Assessing surrounding microglia response to injury**. **(A)** Full field of view representation at 0 minutes and 3 days of neonates with no injury, neonates, adults and aged mice with parenchymal injuries. Asterisks mark where the injury was made. **(B)** Schematic of laser injury experiment. Region of interest is defined by the area outside of the injury ROI. **(C)** Comparison of individual surrouding microglia examples in each experimental group at 0 minites and 3 days after parenchymal injury. Ramification index of surrouding microglia in parenchymal injury experiments. Overall response to parenchymal injury: ****p<0.0001 neonate non-injury and ****p<0.0001 neonate non-injury using Two Way ANOVA and Sidak's post hoc test. Subacute response (pre-injury vs 3 days): ##p=0.0070 neonatal injury vs non-injury; ##p=0.0061 adults vs aged using Two Way ANOVA and Tukey's post hoc test. Data are presented as mean ± SEM. n= 10 fields from 5 P9 non-injury mice, 8 fields from 4 P9 injury mice, n=13 fields from 6 adult mice, n=12 fields from 5 aged mice.
